# Microbes use encapsulin protein organelles to sequester toxic reactions

**DOI:** 10.1101/085266

**Authors:** Tobias W. Giessen, Pamela A. Silver

**Affiliations:** Department of Systems Biology, Harvard Medical School, Boston, Massachusetts 02115, USA; Wyss Institute for Biologically Inspired Engineering, Harvard University, Boston, Massachusetts 02115, USA

## Abstract

Cells organize and regulate their metabolism via membrane-or protein-bound organelles. In this way, incompatible processes can be spatially separated and controlled. In prokaryotes, protein-based compartments are used to sequester harmful reactions and store useful compounds. These protein compartments play key roles in various metabolic and ecological processes ranging from iron homeostasis to carbon fixation. We identified more than 900 proteinaceous encapsulin nanocompartment systems in bacterial and archaeal genomes. Encapsulins can be found in 15 bacterial and 2 archaeal phyla. Our analysis reveals 1 new capsid type and 9 previously unknown cargo proteins targeted to the interior of encapsulins. We experimentally characterize 3 newly identified encapsulin systems and illustrate their involvement in iron mineralization, oxidative and nitrosative stress resistance and anaerobic ammonium oxidation, a process responsible for 30% of the N lost from the oceans. We propose that encapsulins represent a widespread strategy for toxic reaction sequestration in prokaryotes.

---

Cells employ compartmentalization to overcome many difficult metabolic and physiological challenges^1^. Eukaryotes mainly use membrane-bound organelles to sequester and control the flow of metabolites, store genetic information and segregate protein processing and export^2, 3^. In contrast, the majority of prokaryotes do not possess intracytoplasmic membrane systems and instead rely on alternative approaches to achieve spatial control^4, 5^. A number of different protein-based compartmentalization systems have been discovered, most prominently bacterial microcompartments (BMCs) like the CO_2_-fixing carboxysome^6, 7^ and ferritins involved in iron homeostasis and maintaining redox balance^8^. Encapsulating enzymes or biosynthetic pathways in semi-permeable protein organelles can increase the local concentrations of metabolites and enzymes, prevent the loss of toxic or volatile intermediates ^9^ and create unique microenvironments necessary for the proper functioning of specialized enzymes ^10^. In addition, encapsulation allows for incompatible reactions and processes to take place in a single cell at the same time.

A new class of proteinaceous compartments referred to as encapsulin nanocompartments has recently been discovered^11, 12^. They have been reported in a number of bacterial and archaeal genomes and assemble into *T* = 1 (60 subunits, 20-24 nm) or *T* = 3 (180 subunits, 30-32 nm) icosahedral hollow capsids. The key feature of encapsulin systems is that they specifically encapsulate cargo proteins, which are targeted to the encapsulin capsid interior via small C-terminal peptide tags referred to as targeting peptides (TPs)^13^. So far, 7 encapsulin systems have been investigated. Encapsulins internalizing ferritin-like proteins (Flps) were discovered in *Thermotoga maritima*^12^, *Myxococcus xanthus*^14^ and *Rhodospirillum rubrum*^15^ while an Flp-encapsulin fusion protein was found in *Pyrococcus furiosus*^16^. The *M. xanthus* encapsulin system is involved in starvation-induced oxidative stress response, protecting cells from the iron-dependent generation of reactive oxygen species (ROS). A second type of encapsulin system encapsulates DyP-type peroxidases (Peroxi) in *Rhodococcus erythropolis*^17^, *Brevibacterium linens*^18^ and *Mycobacterium tuberculosis*^19^. A role in oxidative stress response was proposed based on cargo protein identity. These examples from diverse microbes raise the question of whether compartmentalization using protein-based organelles is much more widespread than previously assumed.

Here, we systematically explore the distribution, diversity and function of encapsulin gene clusters in bacterial and archaeal genomes. Our approach is the first comprehensive analysis of encapsulin systems in microbial genomes. By surveying and analyzing the newly identified encapsulin gene clusters, we can assess the potential for functional nanocompartment production in organisms occupying diverse habitats ranging from anaerobic hyperthermophilic environments to the human host, and predict their functions at the molecular level. This enables us to formulate informed hypotheses about the roles encapsulin nanocompartments play in survival, pathogenesis and in shaping global ecological processes. We use a combination of bioinformatics, phylogenetic analysis and biochemistry to reveal and characterize 3 new encapsulation systems involved in key cellular processes - iron mineralization, oxidative and nitrosative stress resistance and anaerobic ammonium oxidation (anammox).

## Results

### Encapsulin systems are widespread in bacteria and archaea

The widespread distribution of encapsulin nanocompartments makes them one of the most predominant classes of protein organelles. Our systematic approach identified 909 encapsulin systems distributed in 15 bacterial and 2 archaeal phyla spanning a remarkable breadth of microbial diversity and habitats. Initially, position-specific iterative (PSI)-BLAST searches using all experimentally confirmed encapsulin capsid proteins were carried out against the NCBI database of non-redundant protein sequences. After consolidating the hits resulting from different searches, the 10 kb up- and downstream regions of the identified encapsulin homologs were searched for the presence of phage-related genes using individual protein BLAST searches. When genes related to any known phage morphotype were detected, hits were excluded from subsequent analysis. This step was necessary to ensure that all identified systems are indeed encapsulins and not unknown HK97-related prophages. This resulted in a list of 909 encapsulins distributed in 15 bacterial and 2 archaeal phyla (Fig. 1A and Supplementary Data 1). Encapsulins can be found in anaerobic thermophilic microbes of the genera *Thermotoga, Clostridium* (Bacteria) and *Pyrococcus* (Archaea) as well as aerobic mesophilic soil bacteria (e.g. *Streptomyces* and *Bacillus*), cyanobacteria (e.g. *Acaryochloris*) and host-associated microbes, both commensal (e.g. *Bacteroidetes* and *Propionibacterium*) and pathogenic (e.g. *Mycobacterium, Escherichia* and *Pseudomonas*). Actinobacteria and Proteobacteria harbor the largest number of encapsulin systems (47 and 27%, respectively) while encapsulins can also be found in many less explored phyla like the Planctomycetes and Tectomicrobia. Interestingly, 14 of the identified species contain 2 distinct encapsulin systems (core operons, Data S1). In 8 cases, the systems were of the same core cargo type and likely originated from a duplication of the core operon locus, while in 6 cases, 2 different core cargo types were present (consistently Heme and Peroxi), suggesting independent acquisition via horizontal gene transfer. The presence of multiple encapsulin systems within a single organism argues that each of them conveys a distinct evolutionary benefit.

**Figure 1:**
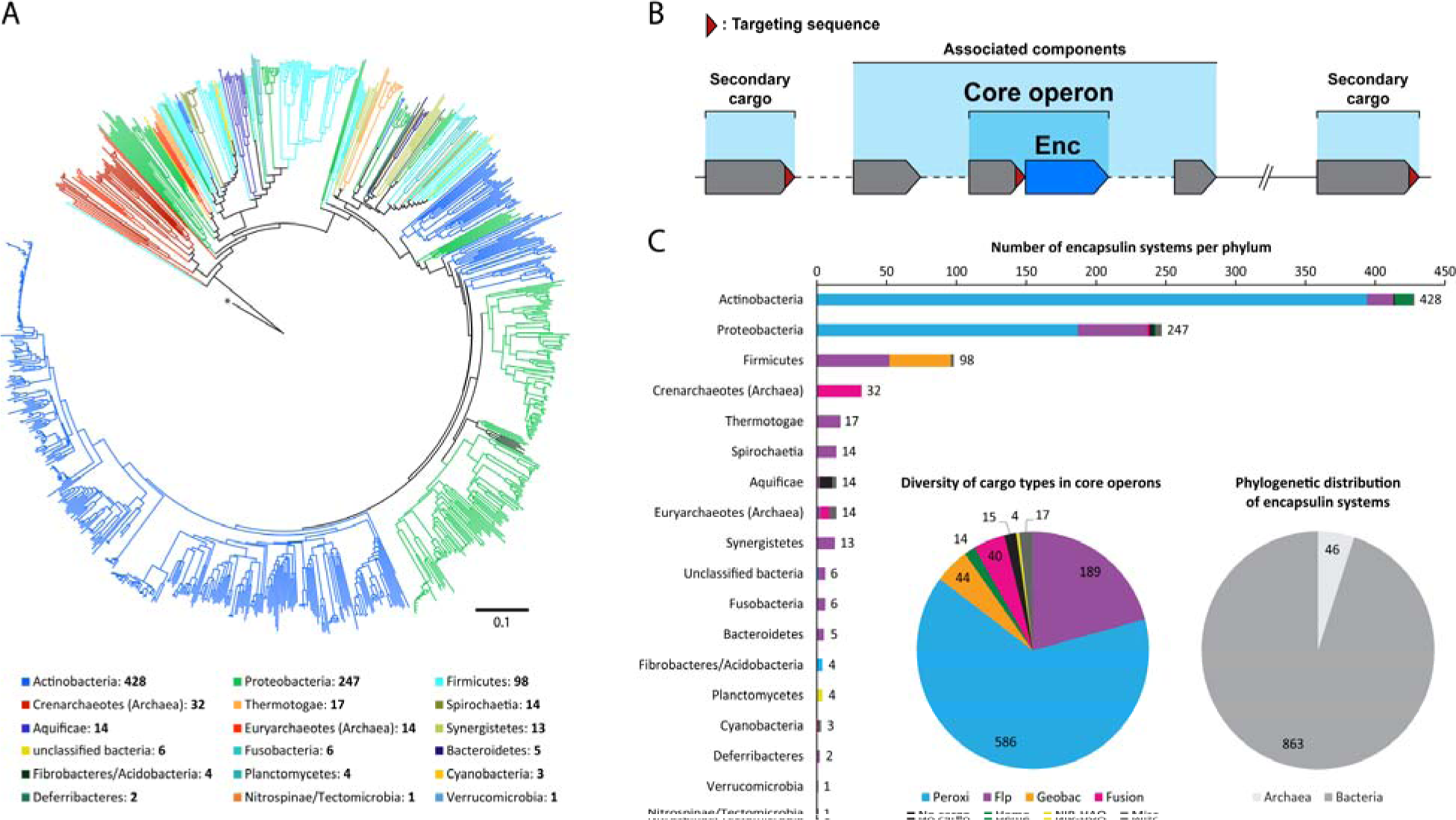
Distribution, organization and diversity of encapsulin systems in prokaryotes. (A) Nearest-neighbor phylogenetic tree of 909 encapsulin capsid proteins using the major capsid protein of bacteriophage HK97 as the outgroup (*) based on a ClustalOmega multiple alignment (distance scale: 0.1). The number of identified encapsulin systems per phylum is given below the phylogenetic tree. (B) Scheme depicting the general organization of encapsulin systems. The core operon consists of an encapsulin capsid gene (Enc) and a core cargo usually directly upstream. Other non-targeted but functionally associated components are found in proximity to of the core operon. Additional secondary cargo can be present throughout the genome. (C) Bar graph depicting the distribution of identified core cargo families in bacterial and archaeal phyla. The number of systems belonging to each identified core cargo type as well as the distribution of encapsulin systems in bacteria and archaea are shown as pie graphs.

### A diversity of cargo types

We identify new cargo types and other encapsulin-associated components illustrating that encapsulin systems are more complex than previously assumed. An encapsulin system is defined by the core operon, composed of the encapsulin capsid and core cargo genes (Fig. 1B). All cargo proteins are characterized by the presence of a targeting peptide (TP) at one of the termini. Additional cargo proteins can be located outside the core operon and are referred to as ‘secondary cargo’. Other associated components that are not targeted for encapsulation, but might play important functional roles can be encoded close to the encapsulin core operon.

Our search reveals 7 previously unknown core cargo types. The set of encapsulin-containing genomes was searched for the presence of core cargo proteins by inspecting the neighborhood of a given encapsulin gene for genes encoding proteins with C- or N-termini similar to known encapsulin TPs using MultiGeneBlast^20^. The 7 newly identified core cargo proteins are generally located directly upstream of a given encapsulin gene (Fig. 1B and C). They include hemerythrins (Heme), di-iron proteins known to coordinate small gaseous compounds like O_2_ and NO^21^, a new type of 4 helix bundle protein (Geobac) prevalent in Firmicutes and 4 distinct Flp-family proteins^22^. In addition, anammox bacteria of the phylum Planctomycetes, capable of carrying out anaerobic ammonium oxidation^23^, exhibit a divergent operon organization with a conserved putative cargo gene located downstream of an unusual fusion-encapsulin. We discovered 4 new secondary cargo proteins and show that up to 4 different cargo types are present in a single genome. One approach to study the physiological functions of encapsulins is to correlate core cargo type with the identity of additional secondary cargo proteins and non-encapsulated genetically associated components^24^. BLAST searches in the set of encapsulin-encoding genomes were carried out using 3 different TP consensus sequences as queries. The first consensus sequence used was based on the TPs of all core cargo types identified in this study. Using the core consensus we identified 76 secondary cargo proteins of 4 different types (Flp: 27, Peroxi: 8, Heme: 14 and Ferredoxin (Fer): 27), 3 of them previously unknown (Peroxi, Heme and Fer) (Fig. 2A). Interestingly, the newly discovered Ferredoxins were present exclusively in Geobac-systems found either directly downstream of the encapsulin gene or separated by one other gene. Additionally, TPs present in Ferredoxins were not located at the C-terminus, as is the case for all other identified cargo, but at the N-terminus. Our search using the FolB-based consensus sequence yielded 76 instances of FolB-type secondary cargo proteins and 8 examples of a new putative cargo type, namely BphB (biphenyl dehydrogenase)-like short chain dehydrogenases. Searches using the BfrB-consensus sequence resulted in the identification of 172 BfrB-type secondary cargo proteins. More than one third (363) of all identified systems contain more than one cargo type that is tagged for encapsulation (Supplementary Fig. 1A). We identified 65 instances of 3 cargo types being present in one genome and one example of 4 co-occurring cargo proteins (core: Peroxi, secondary: Heme, FolB and BfrB; *Mycobacterium simiae* ATCC 25275). Besides core and secondary cargo, non-targeted associated proteins are often found in the vicinity of encapsulin core operons and are confined to specific genera and core cargo types. They include cupin-type oxygenases found in close association with many Peroxi systems in Mycobacteria (Supplementary Fig. 1B), non-targeted Flp proteins found close to Flp or Flp-fusion systems in archaea (Supplementary Fig. 1C) and different types of electron-transfer proteins clustered around Flp core operons (Supplementary Fig. 1D).

**Figure 2:**
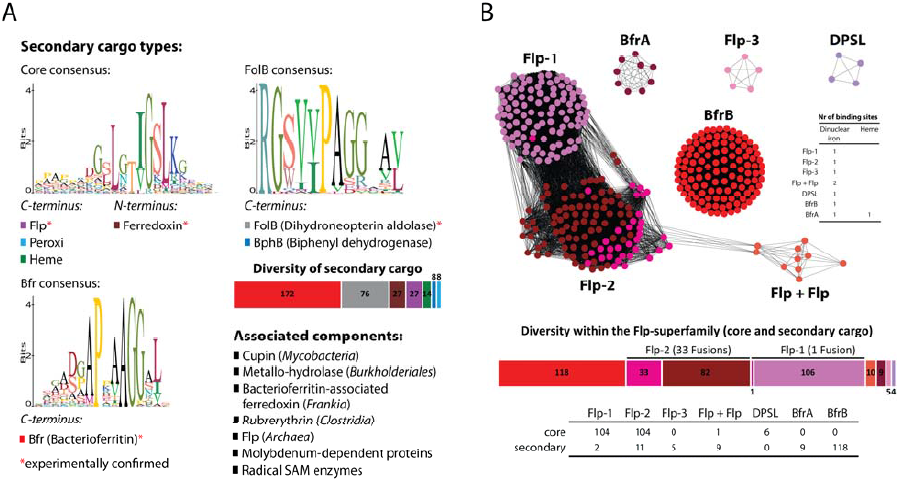
Secondary cargo, associated components and diversity of Flp-like cargo in encapsulin systems. (A) Shown are sequence logos of the 3 targeting peptide consensus sequences used to identify secondary cargo in all encapsulin containing genomes. The y-axis denotes positional information in bits. The secondary cargo types identified using a given consensus as a query are listed below the respective sequence logos. Experimentally confirmed secondary cargo types (either in this study or previously) are indicated with a red asterisk. The number of identified instances of different secondary cargo types are shown as a color-coded bar graph. A list of associated components identified in certain genera (shown in parentheses) is shown. (B) Sequence similarity network of all Flp-family cargo proteins identified in encapsulin systems (edge % identity: 39). The inset table indicates the presence and number of dinuclear iron centers and heme binding sites in each family. A bar graph depicts the number of proteins identified of each Flp-type with Flp-fusions depicted in pink. The bottom table shows the number of sequences of each Flp-family identified as core or secondary cargo.

The most diverse class of identified cargo are Flp-type proteins. They represent a large superfamily with varied functions unified by the presence of a 4 helix bundle domain harboring a dinuclear iron center^22^. A sequence similarity network (SSN) analysis^25^ of all (core, fusion and secondary) Flp-type cargo proteins identified in this study allowed us to visualize distinct subfamilies and showed that 7 distinct classes of Flp proteins are targeted for encapsulation by encapsulin compartments (Fig. 2B). Two (Flp-1 and Flp-2) have previously been identified to be part of encapsulin systems either as distinct cargo proteins (*T. maritima, M. xanthus* and *R. rubrum*)^12, 14, 15^ or as an N-terminal fusion domain (*P. furiosus*)^16^. Flp-3, exclusively identified in Archaea, has not been characterized and shows similarity to proteins found in ferric iron- and arsenic-reducing species^26^. Two types of bacterioferritin-like proteins (BfrA and BfrB) are usually involved in maintaining iron homeostasis in various bacteria representing an analogous system to the eukaryotic Ferritins (Ftn)^22^. Additionally, proteins with similarity to DNA-binding proteins from starved cells (DPSL) were identified in a number of archaeal species. Finally, a previously unknown family of double Flp fusion proteins (Flp+Flp) was shown to be present in a number of thermophilic bacteria (Supplementary Fig. 1E).

### Encapsulins can be divided into four families

Encapsulin nanocompartments were known to assemble into either small (ca. 22 nm in diameter) or large (ca. 32 nm in diameter) capsids with triangulation numbers of *T*=1 (60 subunits) or *T*=3 (180 subunits), respectively^11^. However, it was unclear what factors determine their assembly into small or large compartments or how capsid size correlates with cargo type and function. To address these issues, we initially carried out a SSN analysis of all identified encapsulin capsid proteins (Fig. 3A). We found that encapsulins cluster into 4 distinct groups (edge % identity: 24). It was immediately apparent that all Flp-fusion encapsulins cluster together. By mapping the characterized systems onto our network we found that small (60mer) encapsulins cluster together in one of the 2 large sequence groups while the single known large (180mer) system (*M. xanthus*) was found in the second large sequence cluster (Supplementary Fig. 2A). None of the known systems mapped onto the fourth sequence cluster from now on referred to as ‘New’.

**Figure 3:**
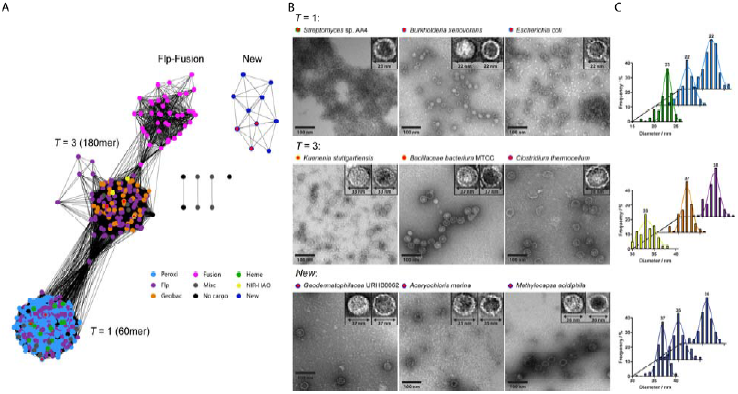
Classification and characterization of encapsulin capsids. (A) Sequence similarity network of all identified encapsulin capsid proteins (edge % identity: 24). Individual nodes are colored corresponding to their associated core cargo type. Red dots indicate the capsids chosen for experimental characterization. (B) Negative-stain TEM micrographs of 9 encapsulin capsids from the *T*=1, *T*=3 and New sequence clusters. Insets show magnified examples of each capsid. (C) Size-distribution of the characterized encapsulin capsids fitted using a Gaussian fit.

There are 2 main determinants at the sequence level predictive of capsid size (Supplementary Fig. 2B). A T3-motif (GPxGxG) present only in large capsids occurs at the N-terminal end of the E-loop while an additional 8 to 10 residue E-loop insertion is present in most *T*=3 capsids. The fourth sequence cluster (New) could not be classified as *T*=1 or *T*=3 due to large sequence insertions and no clear presence of T3-motifs or E-loop insertions. Interestingly, all of the mentioned sequence insertions are present in loops exposed on the outer surface of the encapsulin compartment (Supplementary Fig. 2C). The distinct character of New capsids is additionally supported by the fact that all of them are found in similar gene clusters that are distinct from other encapsulin loci. These clusters often contain genes encoding proteins related to lipid/isoprenoid metabolism and a likely hexameric P-loop NTPase (Supplementary Fig. 3).

All characterized encapsulins were able to assemble into nanocompartments when heterologously produced in *E. coli*, highlighting the inherent robustness and assembly efficiency of encapsulins. Furthermore, these assembled encapsulin nanocompartments could be isolated for biophysical and biochemical analyses. We characterized 3 examples each of the *T*=1, *T*=3 and New clusters. We picked sequences that are diverse both in terms of phylogenetic origin and core cargo type. After heterologous expression and purification (Supplementary Fig. 2D), negative-stain transmission electron microscopy (TEM) was used to visualize encapsulins. As predicted, all *T*=1 type encapsulins assemble into small capsids of 22 (*Burkholderia xenovorans* and *Escherichia coli*) or 23 nm (Streptomyces sp. AA4) in average diameter (Fig. 3B and C). The representatives chosen from the *T*=3 cluster were clearly different and as hypothesized resulted in larger capsids. Interestingly, 2 of the 3 capsids were larger than expected with average diameters of 37 (*Bacillaceae bacterium* MTCC 10057) and 38 nm (*Clostridium thermocellum*) while *Kuenenia stuttgartiensis* capsids were 33 nm in size, similar to the previously reported *M. xanthus* encapsulin capsids (32 nm). The representatives chosen from the New sequence cluster stemmed from 3 different phyla and assembled into larger capsids of 35 (Cyanobacteria: *Acaryochloris marina*), 36 (Proteobacteria: *Methylocapsa acidiphila*) and 37 nm (Actinobacteria: *Geodermatophilaceae* URHB0062), similar in appearance to *T*=3 capsids. Overall, all investigated encapsulin capsids showed very uniform size distributions with standard deviations between 0.9 (*Streptomyces* sp. AA4) and 2.1 nm (*K. stuttgartiensis*) (Fig. 3C).

Certain core cargo types are confined to specific sequence clusters. We mapped the core cargo types of all identified systems onto the SSN (Fig. 3A). Except for Flp-type systems, which were present in both the *T*=1 and *T*=3 sequence clusters, all other core cargo types were confined to a specific capsid type. Peroxi and Heme systems could only be found in the *T*=1 cluster while Geobac and NIR-HAO systems were only present in the *T*=3 cluster. The match of core cargo types with a specific capsid size implies that different capsid sizes have been adapted to serve particular functions.

### A new hemerythrin cargo protects cells from oxidative and nitrosative stress in an encapsulation-dependent manner

Hemerythrins are 4 helix bundle proteins that often form homo- or heterooligomeric complexes with one dinuclear iron center per subunit. In prokaryotes, hemerythrin-like domains have recently been proposed to be involved in O_2_ and NO sensing and in the regulation of oxidative stress response^27^. Hemerythrins, predicted to fold into a 4 helix bundle (Supplementary Fig. 4A), are a new core cargo type (Heme) in encapsulin systems. Heme core operons generally consist of the encapsulin capsid gene and a hemerythrin containing a C-terminal TP encoded directly upstream (Fig. 4A). In one species (*Sporichthya polymorpha* DSM 43042) a second hemerythrin can be found directly downstream of the encapsulin. Most organisms containing a Heme system encode BfrB as an additional secondary cargo at a separate locus in their genomes while in *Pseudonocardia dioxanivorans* CB1190 a Peroxi secondary cargo is present.

**Figure 4:**
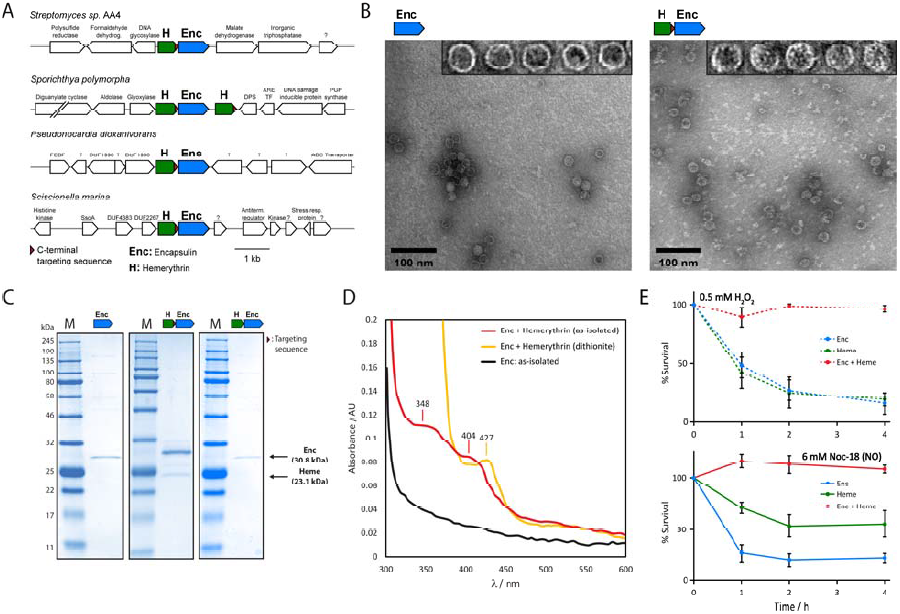
Characterization of the Streptomyces sp. AA4 Heme encapsulin system involved in oxidative and nitrosative stress resistance. (A) Selected Heme encapsulin gene clusters and their genetic surroundings. (B) Negative-stain TEM micrographs of purified encapsulin particles without (left) and with (right) Heme cargo. Insets show magnified particles. Additional protein puncta are visible when both Enc and Heme are present. (C) SDS-PAGE analysis of purified particles without (left) and with (middle) Heme. A mutant Heme missing the C-terminal TP does not co-purify with Enc particles (right). (D) UV-Vis spectroscopy of purified particles. (E) Survival of **E. coli** expressing Enc, Heme or Enc+Heme in the presence of H_2_O_2_ (top panel) or the NO-generator Noc-18 (lower panel). Data are represented as mean ± SD of 3 biological replicates.

Hemerythrin cargo is encapsulated by and co-purifies with encapsulin particles in a TP-dependent manner. We expressed the *Streptomyces* sp. AA4 Heme core operon in *E. coli*. After capsid purification, negative-stain TEM was used for particle visualization (Fig. 4B). Compared with capsids resulting from the expression of the encapsulin gene alone, Heme core operon particles contain additional protein density in the compartment interior. In many cases, cargo was organized in a 5- or 6-fold symmetrical manner appearing as puncta associated with the interior surface of encapsulin particles (Fig. 4B insets and Supplementary Fig. 4B). To confirm that the observed cargo was indeed hemerythrin, we compared SDS-PAGE gels of purified encapsulin and Heme core operon capsids (Fig. 4C). As a control, particles of a system where the C-terminal 12 amino acids of the hemerythrin cargo were removed was used. Only in the wild-type system did hemerythrin co-purify with encapsulin particles confirming the dependence of hemerythrin encapsulation on a C-terminal TP. Based on gel densitometry analysis ca. 20 molecules of hemerythrin were encapsulated per 60 subunit capsid. To investigate the oligomerization state of hemerythrin cargo in the absence of encapsulin we performed analytical size exclusion chromatography (Fig. S4C). We found that hemerythrin cargo was present as a stable dimer in solution.

UV-Vis spectroscopy reveals that the diiron center of encapsulated Heme is protected from strong reducing agents by the encapsulin capsid. Purified Heme core operon capsids showed 2 main absorption peaks at 348 and 404 nm (as-isolated) dependent on the presence of hemerythrin, similar to previously reported spectra of the met-form of hemerythrins where both iron centers are present in their ferric (Fe^3+^) but oxygen-free form (Fig. 4D)^28^. A clear change in the absorption spectrum upon dithionite reduction occurs with the appearance of a new absorption peak at 427 nm. Following dithionite treatment dinuclear iron proteins usually lose all absorption features in this region resulting from the destruction of their metal center. Together, these results indicate that encapsulation protects the metal center from being destroyed by high concentrations of strong reducing agents. Further UV-Vis analysis of non-encapsulated hemerythrin cargo confirmed that without encapsulation all features in the UV-Vis spectrum are lost upon dithionite incubation (Supplementary Fig. 4D).

The Heme containing encapsulin compartment protects *E. coli* from oxidative and nitrosative stress. We carried out experiments investigating the influence of 2 stressors on the survival of *E. coli* expressing the Heme core cargo system, namely hydrogen peroxide (H_2_O_2_) and nitric oxide (NO); hemerythrins detoxify H_2_O_2_^29^ and have been implicated in NO-sensing. Cells expressing either hemerythrin alone, encapsulin alone or the complete Heme core cargo system were exposed to H_2_O_2_ or the NO-generator NOC-18. Samples were taken after different time intervals and plated as serial dilutions for CFU-counting. A strong protective effect against H_2_O_2_ was observed only when both components (encapsulin and hemerythrin) were present while no effect could be detected for hemerythrin alone (Fig. 4E, upper panel). Similarly, the Heme core cargo system was able to completely rescue *E. coli* growth when exposed to NO (Fig. 4E, lower panel). In contrast to H_2_O_2_-exposure, expressing non-encapsulated hemerythrin before applying NO stress led to ca. 50% survival compared with only ca. 20% for the control (only encapsulin capsid present). Additionally, direct interaction of NO with hemerythrin was confirmed via UV-Vis spectroscopy (Supplementary Fig. 4D).

### A new 4 helix bundle protein found in Firmicutes shows encapsulation-dependent iron mineralization and oxidative stress protection

Systematic analysis revealed the presence of a new core cargo type in a subset of Firmicutes annotated as ‘Geobacter domain of unknown function’ (Geobac). In 61% of cases an additional new cargo type, Ferredoxin, is closely associated with Geobac core operons (Fig. 5A). Fer is either located directly downstream of the encapsulin gene, in most cases overlapping with it, or separated by one additional gene whose function is unknown. No other secondary cargo types can be found in genomes encoding Geobac systems. We turned to secondary and 3D structure prediction which revealed that Geobac likely possesses a 4 helix bundle fold with 2 additional shorter C-terminal helices preceding the canonical TP present in all core cargo types (Supplementary Fig. 4A and Fig. 5A). Surprisingly, none of the residues usually associated with iron-coordinating 4 helix bundle proteins are present (Supplementary Fig. 5A).

**Figure 5:**
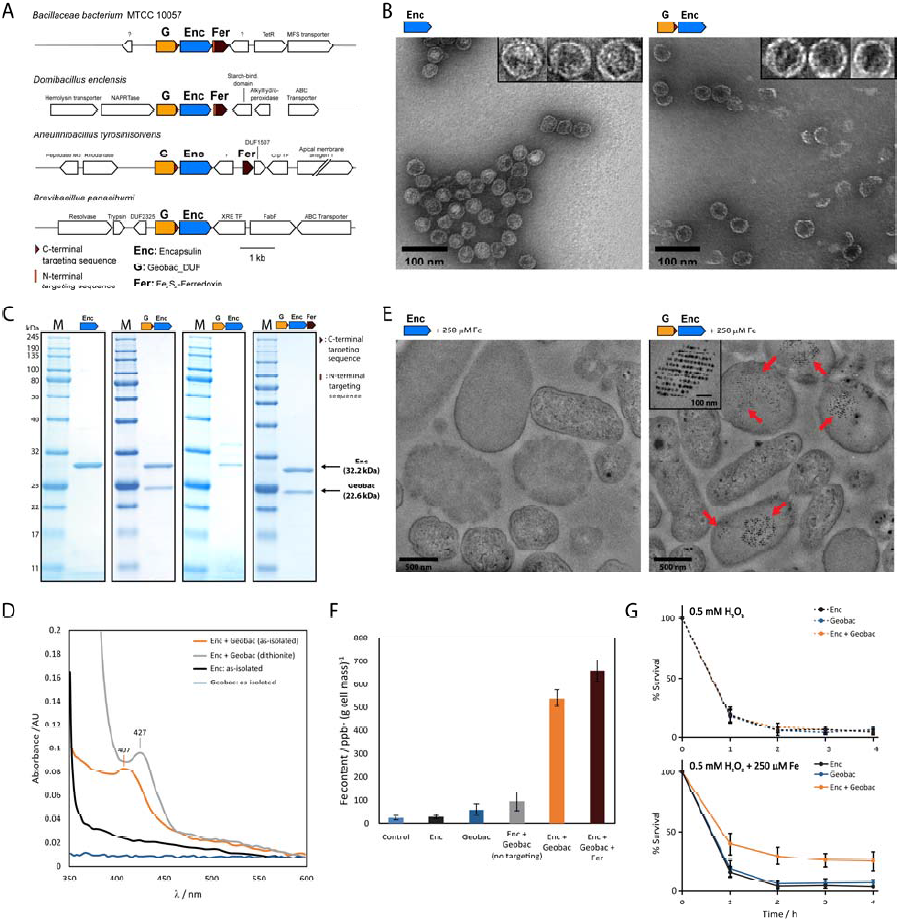
Characterization of the Bacillaceae bacterium MTCC 10057 Geobac encapsulin system involved in iron mineralization. (A) Selection of identified Geobac encapsulin gene clusters. (B) Negative-stain TEM micrographs of purified encapsulin particles resulting from **E. coli** cells expressing Enc alone or the Geobac core operon. Insets show magnified particles. (C) SDS-PAGE analysis of purified particles. The expressed gene/operon is shown above the respective gel. (D) UV-Vis analysis of purified Geobac particles and controls. (E) TEM micrographs of fixed **E. coli** cells expressing Enc or Enc+Geobac in the presence of iron sulfate. Intracellular electron-dense particles are indicated by red arrows. The inset shows a magnification of regularly arranged electron-dense particles found in Enc+Geobac-expressing cells. (F) ICP-MS analysis of washed *E. coli* cells expressing different components of the Geobac encapsulin system. Data are represented as mean ± SD of 3 biological replicates. (G) *Survival of *E. coli** cells expressing Enc, Geobac or Enc+Geobac in the presence of H_2_O_2_ (top panel) or H_2_O_2_ and iron sulfate (lower panel). Data are represented as mean ± SD of 3 biological replicates.

Geobac cargo is encapsulated by and co-purifies with encapsulin particles in a TP-dependent fashion. To further characterize the Geobac core cargo system we focused on the encapsulin, Geobac protein and Fer cargo found in *Bacillaceae bacterium* MTCC 10057. *E. coli*-expression followed by particle purification yielded assembled capsids (Fig. 5B). SDS-PAGE analysis showed the TP-dependent co-purification of Geobac cargo with encapsulin compartments with ca. 150 molecules of Geobac per 180 subunit capsid (Fig. 5C). Analytical size-exclusion chromatography indicated that Geobac cargo exists in an equilibrium between dimer and monomer in solution (Supplementary Fig. 5B). Taken together, the above results suggest that Geobac binds to single encapsulin capsid proteins in a 1:1 or 2:1 ratio, leading to a compartment where the interior surface of the majority of capsid proteins interacts with 1 or 2 cargo molecules.

Fer represents the first encapsulin cargo with an N-terminal TP and co-purifies with encapsulin particles. When expressing the 3-gene operon (Geobac-Enc-Fer), no additional co-purifying cargo could be observed (Fig. 5C) while the 2-gene operon (Enc-Fer) showed a band with the correct apparent molecular weight expected for Fer (Supplementary Fig. 5C and D). Based on gel densitometry, 10 molecules of Fer co-purify per 180 subunit capsid. The identity of this protein band was confirmed to be Fer via in-gel tryptic digest followed by mass spectrometry (Supplementary Fig. 5E). This suggests that the N-terminal TP found in Fer is a weaker targeting signal compared with other C-terminal TPs explaining why initially no co-purifying Fer cargo could be observed.

Geobac is an encapsulation dependent iron binding protein and is able to mineralize iron in vivo. UV-Vis spectroscopy of purified capsids showed an absorption peak at 407 nm only when both Geobac core cargo components were present, which shifted to 427 nm upon dithionite exposure indicating the presence of bound iron (Fig. 5D). Since many 4 helix bundle proteins are involved in iron metabolism ^22^, we carried out experiments where Geobac-expressing cells were exposed to elevated concentrations of iron. Washed cells were either fixed for TEM or subjected to inductively coupled plasma mass spectrometry (ICP-MS). When cells were expressing both Geobac cargo and encapsulin electron-dense puncta were observed while no such puncta where present when only encapsulin was being expressed (Fig. 5E). Electron-dense material was observed in about two thirds of cells and often organized into regular paracrystalline arrangements (Supplementary Fig. 6A and B). The average particle diameter was 23 nm (Supplementary Fig. 6C) suggesting that particles mineralize inside encapsulin compartments (outer diameter: 37 nm, shell thickness: 4-5 nm). ICP-MS analysis was used to compare the iron content of cells expressing different combinations of Geobac system genes. Values adjusted for the protein levels of the Geobac cargo protein (Supplementary Fig. 6D) showed that only when both complete Geobac cargo (with TP) and encapsulin were present could significantly elevated iron levels be observed (Fig. 5F). All other strains showed only background or slightly above background levels of retained iron. Interestingly, cells expressing the 3-gene operon including Fer were able to store the largest amount of iron suggesting that Fer is not necessary for iron mineralization but might improve its efficiency. The Geobac encapsulin system protects *E. coli* from oxidative stress in the presence of elevated levels of iron and likely represents the main iron storage system in Geobac-encoding organisms. We exposed *E. coli* cells expressing Enc, Geobac cargo or both to H_2_O_2_ and monitored the CFU-count over time (Fig. 5G). No increase in survival could be observed when only H_2_O_2_ was present. Only when cells were exposed to both H_2_O_2_ and elevated levels of iron could a clear difference be recorded.

**Figure 6:**
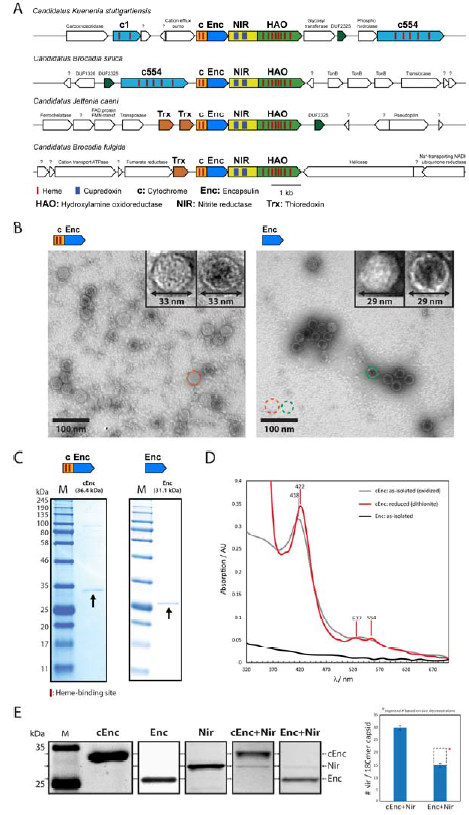
Characterization of the encapsulin system in the anammox bacterium K. stuttgartiensis. (A) Encapsulin operons found in 4 anammox bacteria. (B) TEM micrographs of purified *K. stuttgartiensis* encapsulins with (left) or without (right) N-terminal cytochrome fusion domain. Dotted circles are used to illustrate their size difference. (C) SDS-PAGE gels of cEnc and Enc. (D) UV-Vis spectroscopy of cEnc showing the characteristic absorption pattern of c-type cytochromes. (E) SDS-PAGE gels of purified NIR and different particles from **E. coli** expressing different components of the cEnc+NIR-HAO system (indicated above each lane). The panel on the right shows the decrease in NIR loading when the N-terminal cytochrome domain is missing. Data are represented as mean ± SD of 3 separate purifications and gels.

### Encapsulins in anammox bacteria

Anaerobic ammonium oxidation (anammox, converting NH_4_^+^ and NO_2_^-^ to N_2_) carried out by anammox bacteria of the phylum Planctomycetes was proposed over 50 years ago based on thermodynamic calculations^30^ and is now known to be responsible for over 30% of the nitrogen lost from the oceans^31^. The anammox process is confined to a membrane-bound organelle referred to as the anammoxosome^32^. This organelle is proposed to help sequester highly toxic anammox intermediates (NO, hydrazine and potentially hydroxylamine) thus protecting the rest of the cell^23^. However, it is unclear how the many components inside the anammoxosome are protected from for example NO generated as an intermediate.

We identified encapsulin systems in 4 anammox genomes (*K. stuttgartiensis, B. sinica, J. caeni* and *B. fulgida*, Fig. 6A) encoding a large fusion protein NIR-HAO and an encapsulin N-terminally fused to a cytochrome domain. Anammox encapsulins harbor an additional N-terminal diheme cytochrome domain connected to the encapsulin capsid protein by a ca. 35 amino acid linker rich in proline, alanine and glycine (cEnc, Supplementary Fig. 7A). Based on the presence of CxxCH-motifs we identified the fusion domain as a c-type cytochrome. The 2 heme binding sites both contain additional conserved but different residues (1: CxxxxDCxxCH, 2: ExxxxHxxCxYxxxxCxxCH) that likely influence their coordination sphere and thus redox potential (Supplementary Fig. 7A). The predicted topology of encapsulin capsids results in the N-terminus of capsid proteins protruding toward the interior of the assembled compartment, leading to a particle whose inner surface is lined with diheme cytochrome domains.

In all 4 encapsulin loci identified in anammox bacteria a gene encoding a 2-domain fusion protein (annotated as NIR: nitrite reductase (2 cupredoxin motifs) and HAO: hydroxylamine oxidoreductase (8 heme binding sites)) is located directly downstream of the encapsulin gene (Fig. 6A). Although no TP was present in NIR-HAO, the close association of both genes, their conservation in all 4 genomes and the fact that the associated encapsulin was an unusual fusion protein, led us to hypothesize that NIR-HAO might represent a new type of cargo.

Anammox encapsulins assemble into stable capsids in the presence and absence of their N-terminal c-type cytochrome fusion domain when produced in *E. coli*. We expressed the fusion encapsulin cEnc (*K. stuttgartiensis*) as well as a mutant missing the N-terminal cytochrome domain (Enc) under anaerobic conditions in *E. coli* followed by purification and TEM as well as SDS-PAGE analysis (Fig. 6B and C). Both proteins are able to assemble into stable compartments, but a size difference was observed with Enc forming 29 nm capsids while cEnc formed 33 nm particles. This shows that the N-terminal cytochrome domain influences capsid assembly and thus size. UV-Vis spectroscopy of purified cEnc showed the presence of heme that could be reduced using dithionite resulting in a shift of the 418 nm peak to 422 nm and the appearance of 2 new characteristic peaks at 527 and 554 nm confirming the presence of c-type heme (Fig. 6D).

The NIR domain of the NIR-HAO cargo protein can be produced in *E. coli* under anaerobic conditions and co-purifies with cEnc and Enc. First, we focused on the expression of the 2-gene operon cEnc-NIR-HAO (*K. stuttgartiensis*). However, NIR-HAO was insoluble when expressed under anaerobic conditions in *E. coli*. Difficulties in heterologously producing soluble proteins from anammox bacteria is a well-known phenomenon ^33^. We therefore focused on the N-terminal cupredoxin domain NIR (Supplementary Fig.7C and D). We hypothesized that cargo targeting of the NIR-HAO cargo is mediated by the interaction of NIR and the diheme cytochrome domain of cEnc based on studies showing direct interaction of homologous proteins ^34^. Anaerobically expressing NIR and the 2-gene operons consisting of cEnc/Enc and NIR in *E. coli* indeed yielded soluble NIR (Fig. 6E). NIR co-purified with both cEnc and Enc with ca. 30 (cEnc) and ca. 15 (Enc) copies of NIR per 180mer capsid (Supplementary Fig. 7E). The decrease in the amount of co-purifying NIR for the Enc system cannot be explained by the difference in size between cEnc and Enc alone indicating that the presence of the diheme cytochrome domain improves cargo loading. Based on these results, we propose that NIR-HAO is encapsulated by the fusion encapsulin cEnc.

## Discussion

Here, we systematically identify and analyze nanocompartment gene clusters in bacterial and archaeal genomes and classify them into core cargo groups. We establish a predictive method to organize encapsulin capsid proteins into 4 distinct families, reflecting their size, cargo association and function. We further experimentally characterize 3 newly discovered core cargo systems for assembly, cargo targeting and physiological function establishing their involvement in oxidative and nitrosative stress resistance, iron homeostasis and the anammox process.

The newly identified Heme core cargo system assembles into capsids of the *T*=1 class. The expression of the core operon in *E. coli* confers H_2_O_2_ and NO resistance indicating a protective capacity of the encapsulin system. Previous reports illustrated that hemerythrin model complexes and mutants are able to detoxify H_2_O_2_ via its disproportionation into O_2_ and H_2_O^29, 35^. A similar mechanism in an encapsulation-dependent manner, is likely responsible for the observed increase in survival upon H_2_O_2_ exposure when the complete Heme core cargo system is present. Alternatively, increased survival could stem from efficient sequestration inside the encapsulin compartment or a combination of both active encapsulation-dependent detoxification and sequestration. The partial rescue from NO toxicity observed for cells expressing Heme alone could indicate that resistance is mainly based on NO-binding via hemerythrin and not active detoxification. Encapsulation would then lead to a more efficient sequestration of NO, preventing toxic downstream reactions with O_2_, iron or thiols more thoroughly than free hemerythrin.

The newly identified encapsulin cargo protein Geobac, a 4 helix bundle protein found in Firmicutes, is able to mineralize iron. In 61% of identified Geobac-containing gene clusters, a third gene encoding the new Fer cargo type was identified. This arrangement is reminiscent of bacterioferritin-associated ferredoxins which have been shown to improve mobilization of mineralized iron from bacterioferritins^36^. Fer in Geobac systems could similarly act as an electron-shuttle involved in mineralization or mobilization of iron stored inside the much larger encapsulin compartments. In the presence of elevated iron levels, cells expressing the Geobac core cargo system exhibited improved viability upon H_2_O_2_ exposure. This can be explained by the efficient sequestration of intracellular free iron when the complete Geobac system is present leading to a decrease of toxic ROS-generating Fenton reactions^37^. In addition, iron mineralization associated with ferroxidase activity is known to be able to actively remove H_2_O_2_ in Flp-type proteins^38^. Based on the above results and the fact that all Firmicutes harboring Geobac systems do not encode classical ferritins (Ftn) or bacterioferritins (BfrA or BfrB), we propose that Geobac encapsulin operons represent the main strategy for iron storage in these organism. The fact that 2 conceptually similar but evolutionarily not closely related systems (ferritins and Geobac encapsulins) exist to address the same essential function (iron homeostasis) highlights the power of convergent evolution.

The presence of encapsulins in anammox bacteria is of particular significance given their role in the global nitrogen cycle. Anammox encapsulins are fusion proteins where the encapsulin capsid protein carries a unique N-terminal cytochrome domain. The capsid was shown to assemble into smaller (33 nm) capsids than the other 2 *T*=3 encapsulins analyzed in this study (37 and 38 nm). The N-terminus of encapsulin capsid proteins protrudes towards the interior of the assembled compartment. Thus, the presence of a unique N-terminal domain likely influences capsid assembly and could explain the differing capsid size observed for *K. stuttgartiensis*. All encapsulin loci identified in anammox genomes also possess a conserved TP-less cargo protein NIR-HAO located directly downstream of cEnc. Anammox genomes can contain up to 10 homologs annotated as HAOs (hydroxylamine oxidoreductases) with most of them not having been assigned a physiological function^23^. The HAO domain present in the unique fusion protein NIR-HAO is likely involved in reductive catalysis based on the absence of a heme-coordinating tyrosine residue empirically found to be predictive of oxidative or reductive modes of HAO function^39^. Encapsulation of 3 domains (diheme c-type cytochrome, NIR and HAO) inside a protein compartment suggests that co-localization and sequestration play key parts in this protein organelles’ function. A protein shell would act as a barrier confining toxic intermediates sequestered or generated inside the encapsulin to a dedicated cellular space. Such a system could function in sequestering toxic reactions involved in the anammox process, as a detoxification system inside the anammoxosome and in recycling intermediates lost through diffusion.

The widespread occurrence of protein-based organelles in prokaryotes demonstrates that many bacteria and archaea employ sophisticated compartmentalization mechanisms to spatially organize and regulate their metabolism and adapt to dynamic and hostile environments. Encapsulins are generally found within a subset of a given lineage, often occupying unusual habitats, which argues that they were acquired via horizontal gene transfer rather than vertical inheritance. Encapsulin function is based on the targeted encapsulation of functional protein components and enzymatic functions often involved in carrying out toxic reactions. Thus, we propose that encapsulin systems are a general strategy found in microbes to sequester toxic transformations and molecules and serve as specific storage compartments.

## Methods

### Computational analysis of microbial genomes and encapsulin gene clusters

Genome-mining for encapsulin capsids was initiated by carrying out PSI-BLAST^40^ searches using the following experimentally confirmed capsid proteins as queries: WP_010865203.1 (*T. maritima*), WP_009884820.1 (*B. linens*), WP_020909834.1 (*R. erythropolis*), ABF87797.1 (*M. xanthus*), WP_048059055.1 (*P. furiosus*) and NP_215313.1 (*M. tuberculosis*). Five iterations where run using the following parameters: Expect threshold: 1, word size: 3, matrix: BLOSUM62, gap costs: Existence: 11 Extension: 1, compositional adjustments: conditional compositional score matrix adjustment, PSI-threshold: 0.0005. All hits were combined and the surroundings (10 kb) of the encapsulin gene inspected for the presence of genes annotated as phage-related using BLASTp with default settings. If the case, these hits were deleted. This resulted in a list of 909 encapsulin-encoding loci. Phylogenetic analysis was based on a Clustal Omega (ClustalO)^41^ alignment carried out using the default settings of the Multiple Sequence Alignment online tool of the European Molecular Biology Laboratory’s European Bioinformatics Institute (EMBL-EBI). A nearest-neighbor phylogenetic tree based on the ClustalO alignment was generated using Geneious 9.1.4 with the HK97 bacteriophage capsid protein as the outgroup (NP_037666.1) and then annotated and visualized using the Interactive Tree of Life (iTOL v3) online server^42^.

To identify core cargo proteins present in the set of encapsulin-encoding genomes MultiGeneBlast (MGB)^20^searches using the respective encapsulin proteins and a TP consensus sequence based on characterized core cargo proteins (NP_228595.1 WP_044480584.1 NP_215314.1 WP_009884819.1 ABF88760.1, SDGSLGIGSLKRS) as dual queries were carried out. The consensus was generated using Geneious 9.1.4 and was based on ClustalO alignments. MGB was run in the ‘Architecture search’ mode with minimal sequence coverage at 20%, minimal identitiy at 20%, the maximum distance between genes in locus at 4 kb and the weight of synteny conservation at 1. A custom database of all encapsulin-encoding genomes generated with MGB was used. Only hits where the central residue of the hit string was located closer than 15 residues from the N-or C-terminus were considered valid.

Secondary cargo proteins were discovered using BLASTp searches of encapsulin-encoding genomes with three different TP-consensus sequences as queries. BLASTp parameters were automatically adjusted for short input sequences. The first consensus was generated from all identified core cargo proteins (DGSLGTIGSLKGE). The remaining consensus queries were based on BLASTp hit lists of FolB (RGSVVPAGGAAV) and BfrB (DGAPPAAGGAL) previously identified in *M. tuberculosis*^19^. Alignments were generated using ClustalO and the consensus obtained via Geneious 9.1.4.

Sequence similarity network (SSN) analysis was carried out using the Enzyme Similarity Tool of the Enzyme Function Initiative^25^ with input fasta files containing all protein sequences to be analyzed. After the initial datasets were created we used an alignment score (based on the alignment score vs percent identity plot) that would correspond to a percent identity of 20 for initial outputting and interpretation of the SSN. The resulting xgmml file was imported into Cytoscape 3.3.0. Each node in the network corresponds to a single protein sequence while each edge represents a pairwise connection between two proteins based on a specified % identity cutoff. The cutoffs used for final outputting were 24 (encapsulin capsid proteins) and 39 (Flp-type proteins). The networks were visualized using the yFiles organic layout option of Cytoscape 3.3.0.

### Molecular biology and cloning

All constructs used in this study were ordered as gBlock Gene Fragments from Integrated DNA Technologies (IDT). Codon usage was optimized for **E. coli** with the IDT Codon Optimization Tool with the amino acid sequences of the respective proteins of interest as input. For operons that contained overlapping and/or multiple genes, intergenic regions and overlapping sequences were not changed. Heme, Geobac and NIR cargo constructs for characterization in the absence of encapsulin were ordered containing C-terminal His6 tags. For TP-less Heme and Geobac cargo proteins, the 12 (Heme) and 10 (Geobac) C-terminal residues were omitted, thus removing the TP. The native *K. stuttgartiensis* proteins cEnc and NIR-HAO contain N-terminal signal peptides reminiscent of twin-arginine peptides. They were predicted using the SignalIP 4.1 Server^43^. For cEnc, the N-terminal 36 amino acids (MVMGILNTFKKVYAVTGFFALLAVFSLSQVGSSAFA) and for NIR-HAO the N-terminal 27 amino acids (MLNKSAALVPVVLAFLFLFLCFQCLYA) were omitted when ordering the respective constructs for cloning. The dissected NIR domain of the didomain protein NIR-HAO (CAJ73224.1) was based on the 300 N-terminal residues of the signal peptide-less protein based on domain annotations using Pfam, NCBI-CDD and PROSITE. The N-terminal diheme cytochrome domain of cEnc was removed by omitting the 50 amino acids at the start of the signal peptide-less cEnc protein resulting in Enc. Gibson Assembly^®^ Master Mix was obtained from New England BioLabs (NEB). DNA sequencing was carried out by GENEWIZ. NEB Turbo Competent **E. coli** cells were used for all cloning procedures while BL21 (DE3) Star Competent *E. coli* (NEB) cells were used for protein production and all other experiments. The vectors pETDuet1 and pCDFDuet1 were used for all cloning procedures. For the construction of expression constructs, Gibson Assembly was employed. The respective gBlock Gene Fragments containing 20 bp overlaps for direct assembly were combined with NdeI and PacI digested pETDuet1 or pCDFDuet1 yielding the constructs/strains listed in Table S1. Chemically competent *E. coli* Turbo cells were transformed and the resulting plasmids confirmed by sequencing.

### Expression and purification of proteins and protein compartments

All expression experiments were carried out in lysogeny broth (LB) supplemented with ampicillin (100 μg/mL), spectinomycin (100 μg/mL) or both. Size exclusion chromatography/gelfiltration for capsid purification was performed with an ÄKTA Explorer 10 (GE Healthcare Life Sciences) equipped with a HiPrep 16/60 Sephacryl S-500 HR column (GE Healthcare Life Sciences). For analytical size exclusion to determine oligomerization state in solution a Superdex 200 10/300 GL column (GE Healthcare Life Sciences) was employed. Protein samples were desalted and concentrated with Amicon Ultra Filters from Millipore. For SDS-PAGE analysis, Novex Tris-Glycine Gels (ThermoFisher Scientific) were used. DNA and protein concentrations were measured using a Nanodrop ND-1000 instrument (PEQLab) and UV-Vis spectra were recorded with the same instrument.

Assembled and sequence-confirmed plasmids were used to transform *E. coli* BL21 (DE3) Star cells (ca. 0.5 ng total DNA). For co-transformations of two constructs ca. 15 ng of each plasmid was used. For standard aerobic protein expressions 50 mL LB was inoculated (1:50) using an over-night culture, grown at 37°C and 200 rpm to an OD_600_ of 0.5 and then induced with IPTG (final concentration: 0.1 mM). Cultures were grown at 30°C for 18 h, harvested via centrifugation (4000 rpm, 15 min, 4°C) and the pellets either immediately used or flash frozen in N2(l) and stored at −20°C.

For anaerobic expressions of the *K. stuttgartiensis* constructs cEnc, Enc and NIR, LB medium was inoculated (1:50) and initially grown aerobically to an OD_600_ of 0.5. The culture was then made anoxic through vigorous purging using argon. For all cEnc containing expressions, hemin (stock: 500 μM in 10 mM KOH, final concentration: 10 μM) and freshly made FeSO_4_ (stock: 200 mM in water, final concentration: 10 μM) were added. For NIR expressions CuCl_2_ (stock: 200 mM in water, final concentration: 2 μM) was added. After adding IPTG to a final concentration of 0.1 mM, the cultures were transferred to Vacu-Quik jars (Almore Internationl, inc.). Two BD GasPak EZ satchets were added. This was followed by two rounds of air removal and argon flushing of the jar. Then cultures were incubated at 30°C for 18 h under an argon atmosphere. Cells were centrifuged (4000 rpm, 15 min, 4°C) and the pellets either immediately used or frozen in N_2_(l) and stored at −20°C.

For encapsulin and His-tagged protein purifications, pellets were thawed, resuspended in 5 mL PBS (pH 7.4) buffer, then lysozyme (final concentration: 1 mg/mL) and DNaseI (final concentration: 1 μg/mL) were added and the cells incubated for 20 min on ice. Cell suspensions were subjected to sonication using a 550 Sonic Dismembrator (FisherScientific). Power level 3.5 was used with a pulse time of 8 sec and an interval of 10 sec. Total pulse time was 3 min. Cell debris was subsequently removed through centrifugation (8000 rpm, 15 min, 4°C). The cleared supernatant was then used either for protein affinity or encapsulin particle purification.

His-tagged proteins were purified using Ni-NTA agarose resin (Qiagen) via batch Ni-NTA affinity procedure following the supplier's instructions. Buffer A (50 mM Tris, pH 8, 300 mM NaCl, 2 mM β-mercaptoethanol, 5% glycerol, 20 mM imidazol) was used to wash resin after protein binding and buffer B (50 mM Tris, pH 8, 300 mM NaCl, 2 mM β-mercaptoethanol, 5% glycerol, 250 mM imidazol) was used to elute bound protein. Samples were concentrated and dialyzed using Amicon filters and PBS (pH 7.4) buffer and evaluated using SDS-PAGE. Further analyses were carried out directly or the next day with protein being stored on ice.

To cleared lysates (5 mL) containing encapsulin particles 0.1 g NaCl and 0.5 g of PEG-8000 was added (ca. 10% w/v final concentration), followed by incubation on ice for 30 min. The precipitate was collected through centrifugation (8000 rpm, 15 min, 4°C), suspended in 3 mL PBS buffer and filtered using a 0.2 μm syringe filter. The samples were then subjected to size exclusion chromatography using PBS buffer and a flow rate of 1 mL/min. Fractions were evaluated using SDS-PAGE analysis and protein compartment containing fractions were combined and concentrated using Amicon filters. The samples were either directly subjected to additional analysis or stored on ice overnight. The relative ratios adjusted for molecular mass of encapsulin capsid proteins and co-purifying cargo was determined using the Gel Analyzer of the open source image processing package Fiji based on ImageJ 1.51f.

## Transmission electron microscopy (TEM)

200 Mesh Gold Grids (FCF-200-Au, EMS) were used for all TEM experiments. TEM experiments of fixed cells or negatively stained protein samples were carried out at the HMS Electron Microscopy Facility using a Tecnai G2 Spirit BioTWIN instrument.

For negative-staining TEM, encapsulin samples were diluted to 1-10 μM using Hepes buffer (10 mM Hepes, pH 7.4) and subsequently adsorbed onto formvar/carbon coated gold grids. Grids were glow-discharged using a 100x glow discharge unit (EMS) to increase their hydrophilicity (10 sec, 25 mA) before applying 5 μL of diluted sample. After 1 min adsorption time, excess liquid was blotted off using Whatman #1 filterpaper, washed with H_2_O and floated on a 10 μL drop of staining solution (0.75% uranyl formate in H_2_O) for 35 sec. After removal of excess liquid samples were ready for TEM analysis at 80 keV.

For TEM analysis of fixed cells, 0.5 mL of bacterial culture was fixed by adding fixative (1:1 v/v, 1.25% formaldehyde, 2.5% glutaraldehyde, 0.03% picric acid in 0.1 M sodium cacodylate buffer, pH 7.4). The sample was then incubated at 25°C for 1 h and centrifuged for 3 min at 3000 rpm. The sample containing pellet and supernatant in the same tube was further incubated for 6-18 h at 4°C. Cells were subsequently washed three times in cacodylate buffer, 4 times with maleate buffer pH 5.15 followed by staining with 1% uranyl acetate for 30 min. Then, the sample was dehydrated (15 min 70% ethanol, 15 min 90% ethanol, 2 × 15 min 100% ethanol) and exposed to propyleneoxide for 1 h. For infiltration, a mixture of Epon resin and proylenoxide (1:1) was incubated for 2 h at 25°C before moving it to an embedding mold filled with freshly mixed Epon. The sample was allowed to sink and subsequently moved to a polymerization oven (24 h, 60°C). Ultrathin sections (60-90 nm) were then cut at −120°C using a cryo-diamond knife (Reichert cryo-ultramicrotome) and transferred to formvar/carbon coated grids.

## Determination of average particle diameter

To determine the average diameters of encapsulin particles and electron-dense puncta resulting from Geobac-expression in the presence of elevated levels of iron, TEM micrographs were analyzed using the open source image processing package Fiji based on ImageJ 1.51f. Micrographs were converted to 8-bit binary images, thresholded and processed using the particle analyzer plugin. For encapsulin capsids at least 500 particles were analyzed, while 141 electron-dense puncta (Geobac) were recorded. The diameters reported are based on Fiji Feret diameter output values.

## UV-Vis analysis of Heme, Geobac and cEnc

After purification, as-isolated (oxidized) samples of Heme, Geobac, cEnc and loaded and empty encapsulin capsids were directly subjected to UV-Vis analysis (Nanodrop ND-1000). To study the interaction of Heme, Geobac and cEnc with different small molecule partners, samples were made anoxic through repeated removal of air and flushing with argon using glass vials sealed with rubber septa. Samples in PBS (100 μM) were incubated with sodium dithionite (excess: ca. 50 mM), sodium azide (500 μM), sodium cyanide (500 μM), H_2_O_2_ (500 μM) or diethylenetriamine/nitric oxide (NOC-18, 6 mM) for 5 min on ice before carrying out measurements.

## Oxidative and nitrosative stress survival assays

Overnight cultures of strains expressing different components of the Heme and Geobac encapsulin systems were grown at 37°C and 200 rpm. They were used to inoculate 50 mL LB cultures (1:25) which were grown at 37°C to an OD_600_ of 0.5, then induced using IPTG (0.1 mM) and incubated at 30°C and 200 rpm for 2 h. Cultures were then diluted to an OD_600_ of 0.3 using fresh LB. H_2_O_2_ (final concentration: 0.5 mM) or NOC-18 (final concentration: 6 mM) were then added and cultures incubated at 37°C and 200 rpm in the dark. For Geobac, cultures with and without additional FeSO4 (250 μM) were prepared. Controls did not contain H_2_O_2_/NOC-18. Samples were taken after 0, 1, 2 and 4 h and diluted (1:10-1:1000) using fresh LB. 100 μL of each dilution was then spread on antibiotic-containing LB agar plates which were incubated at 37°C for 18 h. Colonies on plates were manually counted the next day. All experiments were carried out at least in triplicate. Counts were normalized using non-H_2_O_2_/NOC-18 treated cultures and expressed as survival percentages.

## Inductively coupled plasma mass spectrometry (ICP-MS)

ICP-MS measurements were carried out at the Trace Metals Lab at Harvard T.H. Chan School of Public Health. For the analysis of iron content of cells expressing different Geobac components, 1 mL (OD_600_ = 2.4) of induced over-night culture was pelleted and then washed four times with H2O. Pellets were then prepared for ICP-MS analysis using nitric acid resulting in a final concentration of 2% (w/v) and measured with a Perkin Elmer Elan DRC-II Inductively Coupled Plasma-Mass Spectrometer. Reported results are based on three independent measurements per sample.

## In vivo iron mineralization assays

Overnight cultures of cells expressing Geobac constructs were used to inoculate 50 mL LB medium and grown to an OD_600_ of 0.5. Before induction with 0.1 mM IPTG, 250 μM freshly prepared FeSO4 (in water) was added. The cultures were then incubated at 30°C for 18 h. Then cells were fixed for TEM analysis.

## Protein identification

Protein identification of SDS-PAGE gel bands was carried out at the Harvard Medical School Taplin Mass Spectrometry Facility. Coomassie Blue stained gels were destained to clear background. Single bands were excised with as little excess empty gel as possible. Excised gel bands were cut into approximately 1 mm^3^ pieces. Gel pieces were washed and dehydrated with acetonitrile for 10 min. Pieces were then completely dried in a speed-vac. Rehydration of the gel pieces was done with 50 mM NH_4_HCO_3_ solution containing 12.5 ng/μl modified sequencing-grade trypsin (Promega, Madison, WI) at 4°C. After 45 min, excess trypsin solution was removed and replaced with 50 mM NH_4_HCO_3_ solution to just cover the gel pieces. Samples were incubated at 37°C room overnight. Peptides were extracted by removing the NH_4_HCO_3_ solution, followed by one wash with 50% acetonitrile and 1% formic acid. The extracts were then dried in a speed-vac (1 h). The samples were stored at 4°C until analysis. On the day of analysis the samples were reconstituted in 5-10 μL of HPLC solvent A (2.5% acetonitrile, 0.1% formic acid). A nano-scale reverse-phase HPLC capillary column was created by packing 2.6 μm C18 spherical silica beads into a fused silica capillary (100 μm inner diameter, 30 cm length) with a flame-drawn tip. After equilibrating the column each sample was loaded via a Famos auto sampler (LC Packings, San Francisco, CA) onto the column. A gradient was formed and peptides were eluted with increasing concentrations of solvent B (97.5% acetonitrile, 0.1% formic acid). As peptides eluted they were subjected to electrospray ionization and then entered into an LTQ Orbitrap Velos Pro ion-trap mass spectrometer (Thermo Fisher Scientific, Waltham, MA). Peptides were detected, isolated, and fragmented to produce a tandem mass spectrum of specific fragment ions for each peptide. Peptide sequences (and hence protein identity) were determined by matching protein databases with the acquired fragmentation pattern using Sequest (Thermo Fisher Scientific, Waltham, MA). All databases include a reversed version of all the sequences and the data was filtered to between a one and two percent peptide false discovery rate.

## Acknowledgements

This work was supported by a Leopoldina Research Fellowship (LPDS 2014-05) from the German National Academy of Sciences Leopoldina (T.W.G), the DARPA Living Foundries: 1000 Molecules grant (award number: HR0011-14-C-0072; P.A.S) and the Wyss Institute for Biologically Inspired Engineering at Harvard University (T.W.G and P.A.S). We thank Alina Chan and Julian Hegemann for thoughtful comments on the manuscript.

## Author contributions

T.W.G. and P.A.S designed the research and wrote the paper. T.W.G. performed experiments.

